# SARS-CoV-2 lineage B.6 is the major contributor to transmission in Malaysia

**DOI:** 10.1101/2020.08.27.269738

**Authors:** Yoong Min Chong, I-Ching Sam, Jennifer Chong, Maria Kahar Bador, Sasheela Ponnampalavanar, Sharifah Faridah Syed Omar, Adeeba Kamarulzaman, Vijayan Munusamy, Chee Kuan Wong, Fadhil Hadi Jamaluddin, Yoke Fun Chan

## Abstract

**Background:** As of June 30, 2020, Malaysia had confirmed 8,639 cases of COVID-19. About 39% of these were associated with a religious mass gathering event held in Kuala Lumpur between February 27 and March 1, 2020, which drove community transmission during Malaysia’s main wave. We analysed genome sequences of SARS-CoV-2 from Malaysia to understand the molecular epidemiology.

**Methods:** We obtained whole genome sequences of SARS-CoV-2 from 58 COVID-19 patients in Kuala Lumpur, Malaysia, and performed phylogenetic analyses on these and a further 50 Malaysian sequences available in the GISAID database. Malaysian lineage B.6 sequences were further analysed with all available worldwide lineage B.6 sequences.

**Results:** Nine different SARS-CoV-2 lineages (A, B, B.1, B.1.1, B.1.1.1, B.1.36, B.2, B.3 and B.6) were detected in Malaysia. The B.6 lineage was first reported a week after the mass gathering and became predominant (63%) despite being relatively rare (1.4%) among available global sequences. Increases in reported cases and community-acquired B.6 lineage strains were temporally linked. Non-B.6 lineages were mainly associated with travel and showed limited onward transmission. There were also temporally-correlated increases in B.6 sequences in other Southeast Asian countries, India and Australia, linked to participants returning from this event. We also report the presence of a nsp3-C6310A substitution found in 40.5% of global B.6 sequences which has associated with reduced sensitivity in a commercial assay.

**Conclusion:** Lineage B.6 became the predominant cause of community transmission in Malaysia after likely introduction during a religious mass gathering. This event also contributed to spikes of lineage B.6 in other countries in the region.

**Author Summary:** The COVID-19 pandemic in Malaysia was driven mainly by transmission following a religious mass gathering held in Kuala Lumpur at the end of February. To study the genetic epidemiology of SARS-CoV-2 in Malaysia, we analysed 50 available and 58 newly-generated Malaysian whole genome virus sequences. We found that lineage B.6, rare (1.4%) globally, first appeared after the mass gathering and became the most predominant (62.9%) in Malaysia. Increases in COVID-19 cases and locally-acquired B.6 strains were temporally linked. Non-B.6 viruses were mainly associated with travel and showed limited spread. Increases in B.6 viruses in Southeast Asian countries, India and Australia were linked to participants returning from this mass gathering. Altogether, 95.3% of global B.6 sequences originated in Asia or Australia. We also report a mutation in the virus nsP3 gene found in 40.5% of global B.6 sequences and associated with reduced detection by a commercial diagnostic test. In conclusion, the religious mass gathering in Kuala Lumpur was associated with the main wave of COVID-19 cases of predominantly B.6 lineage in Malaysia, and subsequent spread of B.6 viruses regionally. Genome sequence data provides valuable insight into virus spread and is important for monitoring continued accuracy of diagnostic kits.

## Introduction

Coronavirus disease (COVID-19), caused by severe acute respiratory syndrome coronavirus 2 (SARS-CoV-2), has caused more than 20 million infections and 746,000 deaths globally [1]. In Malaysia, there have been 8,639 confirmed cases with 121 deaths reported as of June 30, 2020 [2]. COVID-19 was first identified in Malaysia on 25 January 2020 with early sporadic cases mainly associated with travel from China and Singapore. The main wave of infection began in early March 2020, and was associated with attendance at a mass religious gathering held by the Tablighi Jamaat group in Kuala Lumpur between 27 February to 1 March. This gathering was attended by 16,000 people, including about 1,500 from overseas, including Australia, Canada, Nigeria, and India, and neighbouring Southeast Asian countries Singapore, Indonesia, Thailand, Cambodia, Vietnam, Brunei and Philippines [3]. As the number of confirmed cases rapidly increased, Malaysia had the highest number of confirmed cases in Southeast Asia in March and early April. A movement control order, or lockdown, restricted movement except for necessity, work, and health circumstances from 18 March. Through rigorous public health measures, case numbers began to fall, allowing phased lifting of restrictions after 3 months.

SARS-CoV-2 is an RNA virus from the family of *Coronaviridae* and genus *Betacoronavirus.* There are currently more than 80,000 publicly available complete or near-complete genome sequences of SARS-CoV-2 with over 100 lineages identified. As the virus is rapidly evolving, it is important to understand the genomic epidemiology of SARS-CoV-2 to track virus evolution and local and global spread.

Here we report the whole genome sequences of SARS-CoV-2 from 58 patients from University Malaysia Medical Centre (UMMC), a designated COVID-19 hospital in Kuala Lumpur. Our objective was to study the genetic diversity and epidemiology of SARS-CoV-2 in Malaysia, relative to the regional and worldwide virus lineages. In particular, we were interested in determining the role of the Tablighi Jamaat gathering in regional spread.

## Methodology

Patients admitted to UMMC with suspected COVID-19 were diagnosed by real-time PCR detection of SARS-CoV-2 in nasopharyngeal and oropharyngeal swabs, using a WHO-recommended Berlin Charité protocol [4] and commercial assays. For each patient, the earliest available sample during illness with the highest viral load was selected for direct sequencing. Cases were diagnosed between February and April 2020. This study was approved by the University Malaya Medical Centre ethics committee (no. 2020730-8928).

Viral RNA was extracted from 58 primary samples using QIAamp Viral RNA Mini Kit (Qiagen, Germany) and amplified according to the ARTIC-nCoV-2019 protocol [5]. Briefly, cDNA was synthesized using SuperScript IV First-Strand Synthesis System (Invitrogen, USA) with random hexamers. Multiplex PCR was performed with Q5 High-Fidelity DNA Polymerase (NEB, USA) using two pools of 109 nCoV-2019/V3 primer sets to generate overlapping 400 nucleotide amplicons. PCR products were pooled into one tube and purified using AMPure XP beads (Beckman Coulter, USA). The iTrue library method was used for library preparation [6]. Libraries were sequenced using iSeq 100 reagent kit (Illumina, USA) on the iSeq 100 system (Illumina, USA), with output of 1 × 300bp reads.

The sequenced reads were analysed using Geneious Prime 2020 (Biomatters, New Zealand). Reads were trimmed for quality using default parameters and mapped to reference strain Wuhan-Hu-1 (GenBank accession number MN908947). Leftover gaps were sequenced separately by conventional Sanger sequencing using the nCoV-2019/V3 primers set and consensus sequences were generated. Multiple sequence alignment was performed using MAFFT with default parameters [7]. Phylogenetic analysis was conducted with RAxML 8.2.11 implemented in Geneious with default parameters (generalized time-reversal (GTR) + gamma substitution model and bootstrapped 1000 times) using sequenced samples and 50 other Malaysian genome sequences available at GISAID (www.gisaid.org) as of July 17 2020 [8]. Variant analysis was conducted among all Malaysian sequences using Geneious. Nucleotide mutations were further confirmed using CoV-GLUE online tool (http://cov-glue.cvr.gla.ac.uk) [9]. Lineage groups were classified according to the Pangolin COVID-19 Lineage Assigner online tool (www.pangolin.cog-uk.io) [10].

SARS-CoV-2 lineage B.6 complete genome sequences up to 17 July 2020 were retrieved from GISAID. A final dataset of 928 sequences were aligned with the 42 new B.6 sequences generated in this study using MAFFT with default parameters in Geneious. The alignment was then subjected to maximum-likelihood (ML) phylogenetic analyses using IQTREE v1.6.12 [11], using the GTR+F+I+G4 nucleotide substitution model and assessing branch support by the Shimodaira-Hasegawa-like approximate likelihood ratio test with 1,000 replicates. The ML tree was visualized using FigTree v1.4.

## Results

Fifty-eight whole genome sequences with >99% reads mapped to the reference genome were generated, with average coverage depth of 944× (range, 74× to 7119×; S1 Table) The consensus sequences have been deposited in the GISAID database (accession numbers EPI_ISL_501176 to EPI_ISL_501228, and EPI_ISL_506996 to EPI_ISL_50700). All other Malaysian whole genome sequences (50 sequences) available in GISAID were added to the analysis, making a total of 108 Malaysian sequences.

Nine lineage groups were identified in Malaysian sequences: A, B, B.1, B.1.1, B.1.1.1, B.1.36, B.2, B.3 and B.6 (Fig 1). The most identified lineage group was B.6 (n=68, 62.9%), followed by B (n=22, 20.4%), B.1.1 (n=5, 4.6%), B.1 (n=4, 3.7%), A (n=3, 2.8%), B.1.1.1 (n=2, 1.9%), B.2 (n=2, 1.9%), B.1.36 (n=1, 0.9%) and B.3 (n=1, 0.9%).

**Fig 1.**
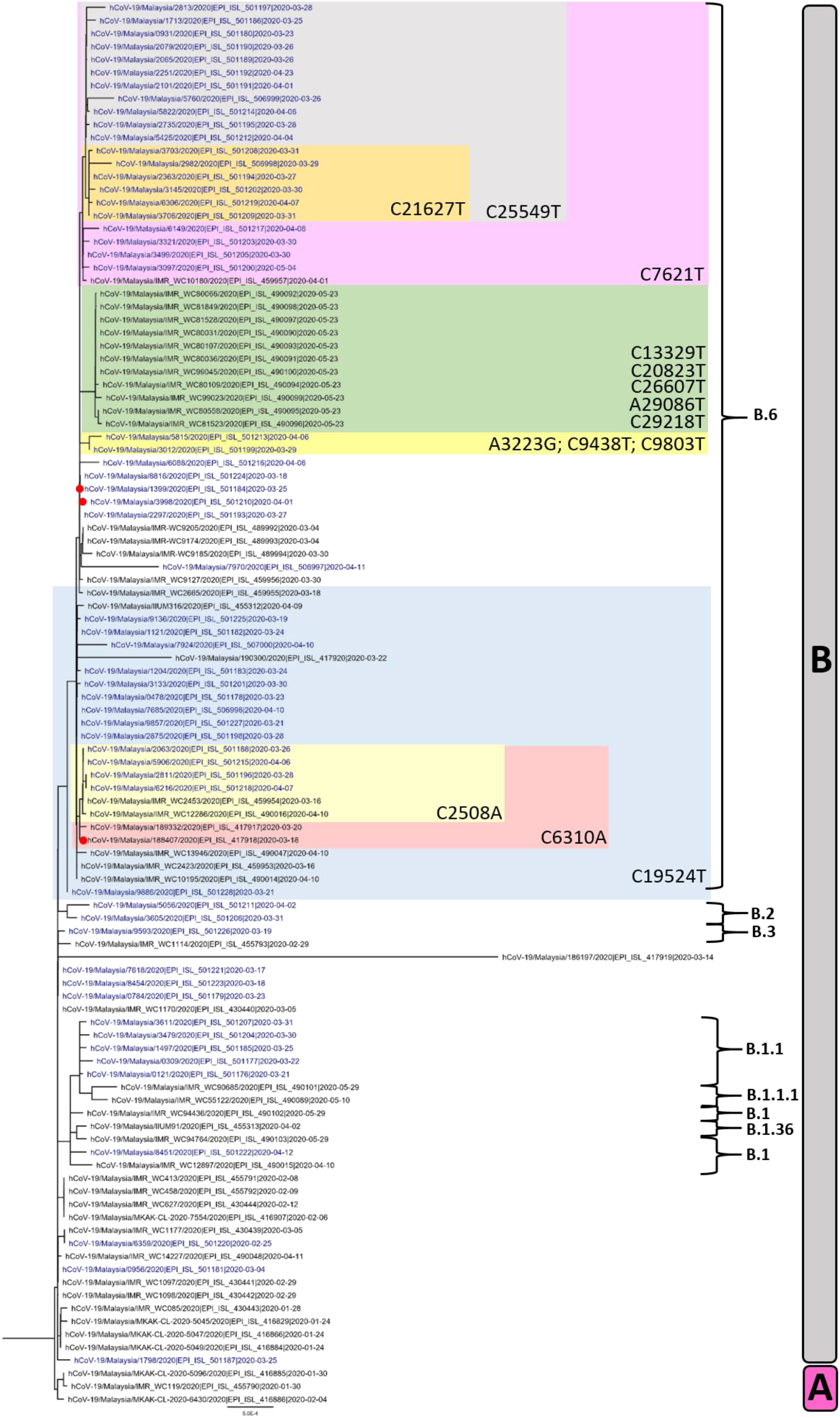
Phylogenetic tree of 108 SARS-CoV-2 whole genome sequences from Malaysia. Sequences from our study are shown in blue and 3 sequences linked to the Tablighi Jamaat gathering are denoted with red circles. The tree is rooted on the branch separating lineages A and B. The key single nucleotide variants in lineage B.6 are highlighted.

Although lineage B.6 was the most common in Malaysia, only 970 B.6 sequences (including those in this study) had been reported globally out of over 60,000 sequences available as of 17 July 2020. Seventeen of the sequences generated from the current study were part of a large healthcare-associated cluster, but B.6 remained the most frequently detected lineage. Lineage B.6 sequences were present in 42/57 (73.7%) of study sequences and 26/39 (66.7%) of other Malaysian sequences in GISAID dating from the Tablighi Jamaat gathering, indicating that this finding was not just confined to our centre.

Among the 68 Malaysian B.6 sequences, a few distinct mutations were observed. The 17 sequences in the healthcare-associated cluster had a non-synonymous mutation C25549T (P53F) in ORF3a (Fig 1). A secondary cluster within this healthcare-associated cluster contained six sequences with an additional mutation C21627T (T22I) in the spike protein. Eight Malaysian sequences, including 6 from this study, had the additional substitution C6310A, resulting in the amino acid change S1197R in nsp3. This C6310A mutation led to reduced sensitivity by 1,000-10,000 viral copies (>10 cycles) with one of the commercial real-time PCR assays used in our laboratory when compared to the well-established Charité assay [4] and updated primers/probes issued by the assay manufacturer in response to this (S2 Table). Some mutations were unique to Malaysian B.6 genomes, such as C2508A, A3223G, C9438T, C9803T, C13329T, C20823T, C26607T, A29086T and C29218T; others, including C6310A, T7621C and C19524T, were also observed in B.6 sequences from other countries including Singapore, Australia and India (Fig 2).

**Fig 2.**
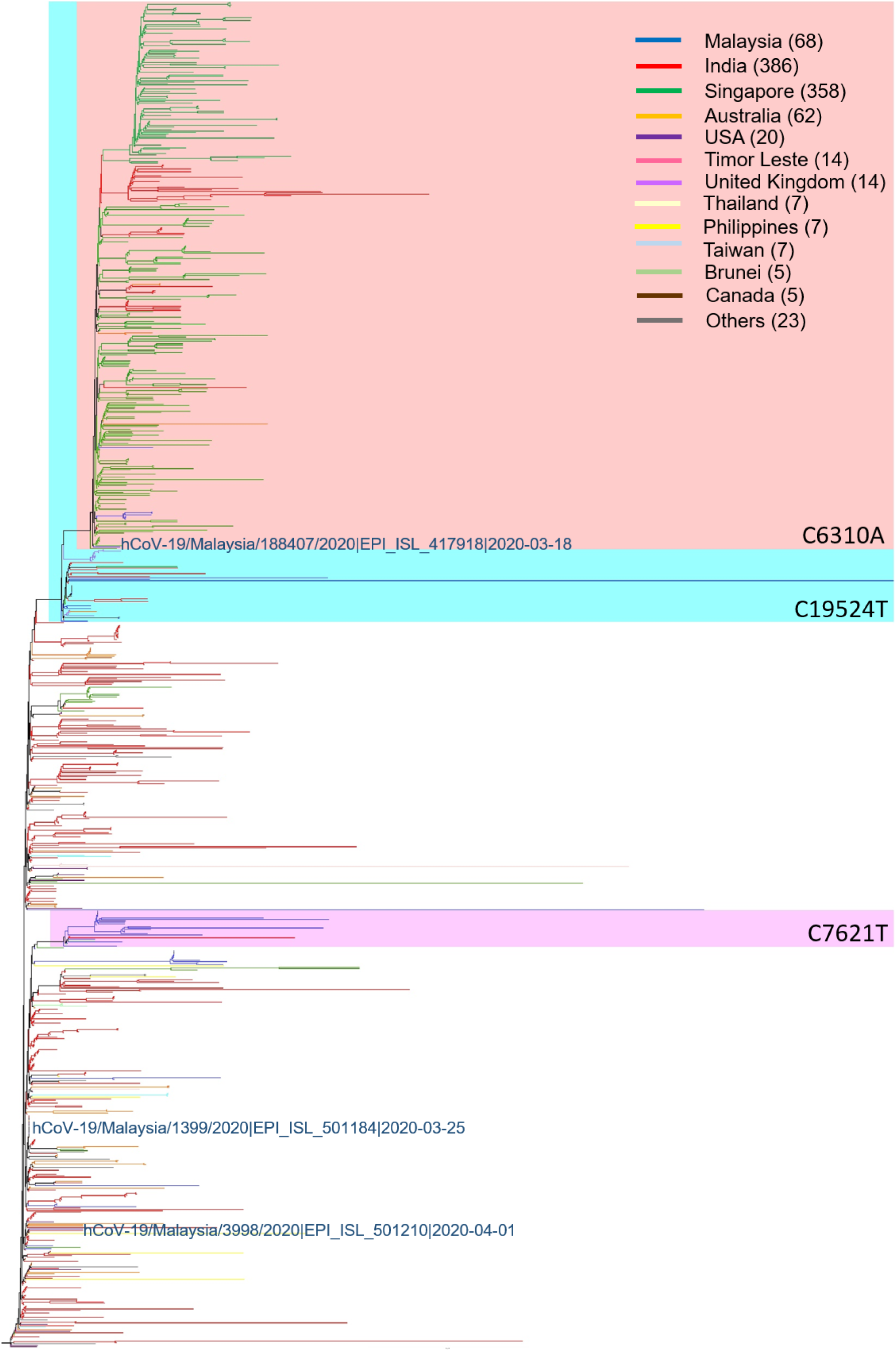
Phylogenetic tree of all 970 B.6 lineage sequences available from the GISAID database as of 17 July 2020. Sequences linked to the Tablighi Jamaat gathering are labelled in blue.

The first reported B.6 sequences globally were from Malaysia and Taiwan on March 4, and the spike of lineage B.6 in Malaysia and other countries clearly began about a week after the Tablighi Jamaat ended on March 1 (Fig 3A and 3B). The 970 reported B.6 sequences represented just 1.4% of the sequences in GISAID, with 924 (95.3%) originating from Asia or Australia. India had the highest number of sequences (386), followed by Singapore (358), Malaysia (68), Australia (62), USA (20), Timor Leste (14), United Kingdom (8), Thailand (7), Taiwan (7), Philippines (7), Brunei (5) and Canada (5). In several Southeast Asian countries (Malaysia, Singapore, Brunei and the Philippines), lineage B.6 sequences accounted for 30.4% −100% of total sequences in GISAID (Fig 3C). At least 3 of the Malaysian B.6 cases in our current and previous studies [12] had direct epidemiological links to the Tablighi Jamaat gathering. Of these, sequence 188407 clustered with sequences from Singapore, India and Australia; sequence 1399 clustered with sequences from India and the Philippines; and sequence 3998 clustered with those from India, USA, Oman, Australia, Taiwan, Canada and Guam (Fig 2).

**Fig 3.**
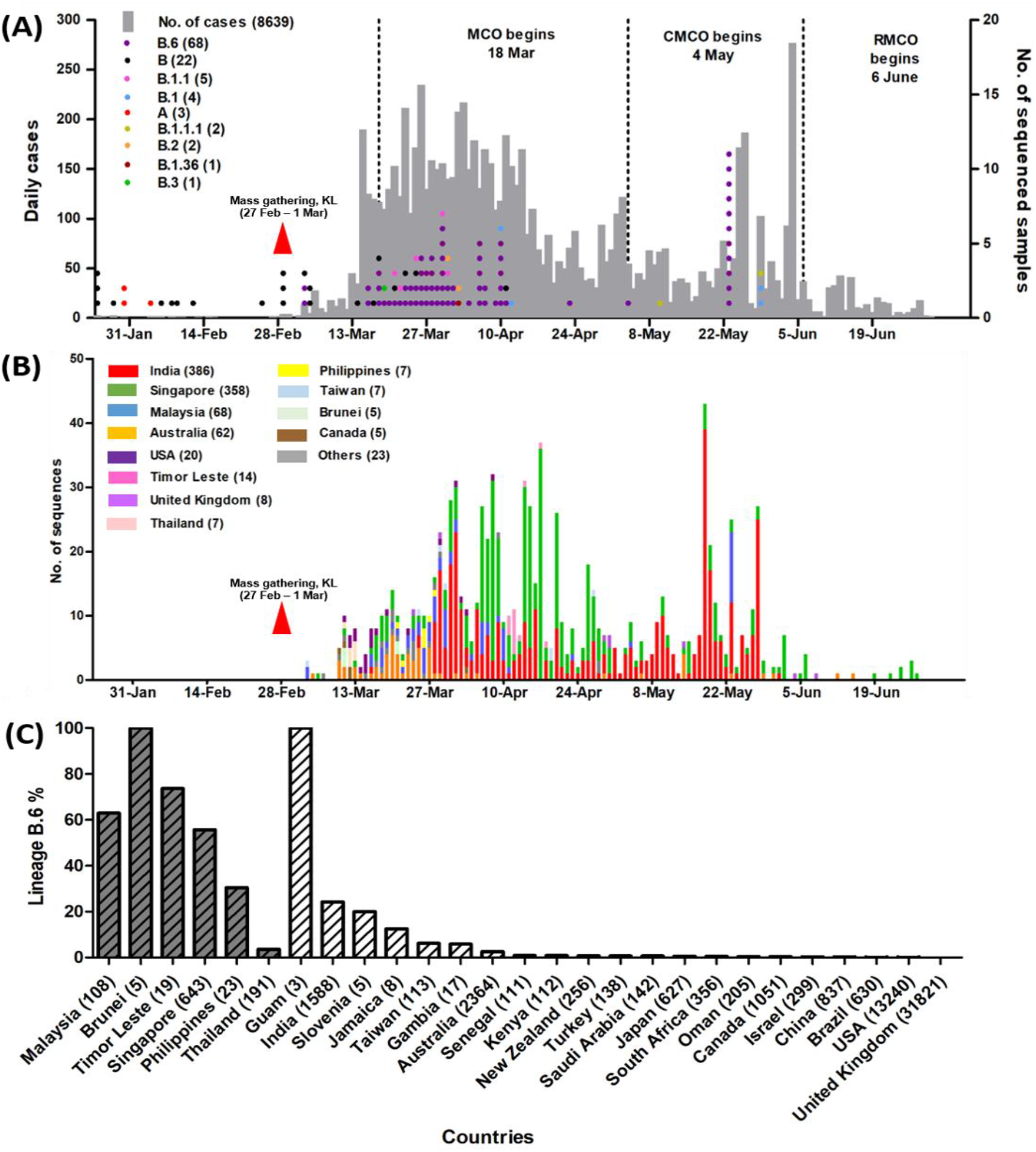
**(A) Epidemic curve of COVID-19 in Malaysia and time course of reported lineages based on 108 Malaysian sequences, from the first reported case on 25 January to 30 June 2020.** The Tablighi Jamaat gathering in Kuala Lumpur is labelled as “mass gathering, KL”. The different phases of lockdown are shown, comprising the movement control order (MCO), conditional movement control order (CMCO) and recovery movement control order (RMCO). **(B) Time course of reported lineage B.6 sequences available in the GISAID database (as of 17 July 2020).** Countries with fewer than 5 lineage B.6 sequences are included in “others”, comprising Brazil (1), China (2), Gambia (1), Guam (3), Israel (1), Jamaica (1), Japan (4), Kenya (1), New Zealand (2), Oman (1), Saudi Arabia (1), Senegal (1), South Africa (2), Slovenia (1) and Turkey (1). **(C) The rates of lineage B.6 sequences over total available complete sequences from each country available at GISAID as of 17 July 2020 (in brackets).** Southeast Asian countries are colored dark grey.

## Discussion

In Malaysia, the Tablighi Jamaat mass gathering resulted in the screening of over 42,000 individuals, with 3,375 confirmed cases (39% of national cases) and 34 deaths, over multiple generations and at least 17 sub-clusters. Of the cases, 825 (24.4%) were foreign nationals from 28 countries, including Southeast Asian countries, Australia, India, Pakistan and Bangladesh [13]. Religious gatherings were recognized early in the pandemic as a significant factor in the spread of COVID-19 [14], with large clusters associated with events in India, Iran and South Korea [15–18].

On March 4, a week after the Tablighi Jamaat event started, two sequences from Malaysia were the first of the B.6 lineage to be reported globally. The diversity of 3 sequences with known epidemiology links to the event (Fig 2) suggests there may have been multiple strains involved in the initial exposure, which is compatible with the large number of attendees from many different countries. The Tablighi Jamaat-associated rise in COVID-19 cases and temporally-correlated rise in reported B.6 sequences in Malaysia, as well as B.6 in other Southeast Asian countries and Australia and India support the role played by local and foreign participants at this event. Attendees at the Kuala Lumpur event also apparently acted as sources of further extensive transmission occurring at two subsequent Tablighi Jamaat gatherings in Pakistan and India later in March [19]. This may explain the later rise in B.6 in India in late March (Fig 3B), such that India now reports the highest number of lineage B.6 sequences as of 19 June. The first known COVID-19 case in Brunei returned from the Kuala Lumpur mass gathering and 52.6% of subsequently reported Brunei cases were associated with this event as of early April [20]. Southern Thailand also experienced an outbreak of COVID-19 after Thai participants returning from this event had a reported 29.3% attack rate [21]. Other Southeast Asian countries including Cambodia, Indonesia, Singapore, Philippines and Vietnam also reported cases associated with this gathering [22–26]. A caveat is that the number of whole genome sequences reported from developing countries in Asia is relatively low and likely underreports the true incidence of B.6. It is notable that this lineage is a very minor contributor in developed countries with greater sequencing volumes (Fig 3C).

Whole genome sequencing of SARS-CoV-2 in our centre not only allowed important insights into local epidemiology, but was also used to track networks within a healthcare-associated cluster, which will be reported elsewhere. Furthermore, public sharing of sequences contributed to identification of the C6310A (nsp3-S1197R) mutation affecting the sensitivity of a commercial PCR assay, leading to updated primers/probes. This mutation is present in only 0.8% of global sequences, but as many as 393 (40.5%) of 970 available lineage B.6 sequences, and within our centre we estimate that the reduced sensitivity of the original assay impacted 10% of positive samples. With many diagnostic kits now on the market and the continued evolution of the virus, this shows the importance of choosing reputable manufacturers who diligently monitor new circulating sequences for potential primer/probe mismatches [27].

There were 40 (37.0%) non-B.6 lineage sequences reported from Malaysia. At least 28 (70%) reported recent international travel (including 3 of the earliest cases from lineage A, imported from China), compared to 2/42 (4.8%) of our B.6 cases. Our sequenced cases with lineages B.1.1, B.2 and B.3 had travelled from Europe and a single B. 1 case had returned from a cruise to the Americas. The B.1 and B.1.1 lineages are the most common in North America and Europe [28]. The mutation D614G in the spike protein, which may increase infectivity of SARS-CoV-2 [29] and has become prevalent in many countries, was only observed in 12 (11.1%) Malaysian sequences from lineages B.1, B.1.1, B.1.1.1 and B.1.36 (S3 Table), but was not present in lineage B.6. The small numbers of other non-B.6 lineages indicate multiple introductions from overseas that failed to establish significant transmission in Malaysia, as the movement control order from 18 March closed the country’s borders and mandated quarantine for all incoming travellers. However, by this point the B.6 lineage had established community transmission, reflected by the local acquisition of disease in most of our B.6 cases without direct or discernible links to the Tablighi Jamaat event.

In summary, in Malaysia COVID-19 spread and the predominance of lineage B.6 was associated with the Tablighi Jamaat religious gathering. Attendees at this event likely spread strains of the B.6 lineage to other countries in the region, including Southeast Asian countries, India and Australia. Further sequence and epidemiological data is needed from Malaysia and other countries in the region to understand the full extent of the spread of B.6. The use of genomic data is important to rapidly identify possible transmission chains and provide a framework for the response to COVID-19.

## Supporting Information

**S1 Table. List of 108 SARS-CoV-2 genomes derived from Malaysian samples used in this study.** The first 58 sequences were generated in this study.

**S2 Table. Comparison of threshold cycle (Ct) values obtained using the Berlin Charité RT-qPCR assay (E gene) [4] and commercial RT-qPCR assay (nsp3 gene; original and updated primers/probes) using 16 patient samples from this or a previous study [12] categorised by presence/absence of the nsp3-C6310A substitution.** A difference in viral RNA of one log or 10× in the initial template concentration is equivalent to a Ct difference of 3.3.

**S3 Table. Nucleotide and amino acid changes in different SARS-CoV-2 lineages in the 108 Malaysian sequences compared to the reference strain Wuhan-Hu-1 (MN908947).** Amino acids are denoted in parentheses and unique mutations found in each lineage are bold and highlighted in grey.

## Acknowledgements

This study was funded by the Defense Threat Reduction Agency, USA under Broad Agency Announcement HDTRA1-6 (grant number HDTRA1-17-1-0027). The funder had no role in the study or publication. We gratefully acknowledge the authors from originating and submitting laboratories of GISAID sequence data on which the analysis is based. We thank GenSeq Sdn Bhd, Malaysia for their assistance in sequencing. The authors are part of the University Malaya COVID-19 Research Group, which include the healthcare workers involved in care of COVID-19 patients in the University of Malaya Medical Centre.

